# Visualizing movements in *E. coli* F_1_F_o_ ATP synthase indicates how the F_1_ and F_o_ motors are coupled

**DOI:** 10.1101/622084

**Authors:** Meghna Sobti, James L. Walshe, Robert Ishmukhametov, Alastair G. Stewart

**Affiliations:** St Vincent’s Clinical School, Faculty of Medicine, UNSW Sydney, Kensington, NSW 2052, Australia; Molecular, Structural and Computational Biology Division, The Victor Chang Cardiac Research Institute, Darlinghurst, NSW 2010, Australia; Department of Physics, Clarendon Laboratory, University of Oxford, Oxford, United Kingdom

## Abstract

F_1_F_o_ ATP synthase functions as a biological rotary generator and makes a major contribution to cellular energy production. It is comprised of two motors that are coupled together by a central “rotor” and peripheral “stator” stalk. Proton flow through the F_o_ motor generates rotation of the central stalk that induces conformation changes that catalyze production of ATP in the F_1_ motor. Here we provide 3-4 Å resolution cryo-EM structures of *E. coli* F_1_F_o_ ATP synthase in 10 mM MgADP. In addition to generating a comprehensive structural model of *E. coli* F_1_F_o_ ATP synthase to provide a framework to interpret mutagenesis studies, we describe a rotational sub-step of the F_o_ motor *c*-ring associated with long-range conformational changes that suggests an elegant mechanism by which the F_1_ and F_o_ motors can be coupled with minimal energy loss.

## Main Text

A key component in the generation of cellular metabolic energy is F_1_F_o_ ATP synthase, a biological rotary motor that converts the proton motive force (pmf) to adenosine tri-phosphate (ATP) in both oxidative phosphorylation and photophosphorylation (*1–3*). The enzyme is comprised of two rotary motors, termed F_1_ and F_o_, that are coupled together by two stalks; a central “rotor” stalk and a peripheral “stator” stalk (*4*), with the simplest subunit composition found in bacteria such as *Escherichia coli*. The F_o_ motor spans the membrane and converts the potential energy from the pmf into mechanical rotation of the central rotor that, in turn, drives conformational changes in the catalytic F_1_ motor subunits to generate ATP (*5, 6*). The F_o_ motor has been hypothesized to act as a ‘Brownian ratchet’, whereby thermal fluctuations and sequential binding of protons to a ring of subunits in the rotor drives mechanical stepping of the central stalk rotor, akin to a water wheel (*7–9*). In *E. coli* the F_o_ motor contains ten of these subunits (*c*_10_) arranged in a ring and so the rotor rotates 36° for each proton that binds (*10*). Conversely, the catalytic F_1_ motor contains a hexamer of subunits (α_3_β_3_) containing three catalytic sites, with one ATP molecule produced with each 120° movement of the rotor (*5, 6*). Because there is a symmetry mismatch between the F_1_ and F_o_ motors that results in a nonintegral H+/ATP ratio (*11*), the coupling between the two motors must be dynamic to enable the enzyme to operate smoothly and function with high efficiency. The nature of this flexibility and the contribution made by the stator and rotor has been debated extensively, with hypothesises proposing flexibility in regions within either the central (*12, 13*) or peripheral stalks (*14*). In a recent cryo-EM study of *Polytomella* mitochondrial ATP synthase, flexible coupling was mainly observed in the *OSCP* subunit that binds the peripheral stalk to the complex at the top of the F_1_ motor rather than to the stalks themselves (*15*).

To investigate the origin of this flexible coupling and rotor/stator interface in ATP synthases more generally and to provide insight into the intermediate steps in the enzyme cycle, we used cryo-EM to probe the structure of detergent solubilized cysteine-free *E. coli* ATP synthase, a widely-used model system for ATP synthases (Fig. 1, S1 and S2). The maps obtained generated far superior structural information than was observed previously for this complex (*16, 17*), with resolutions calculated to be in the range of 3-4 Å. Moreover, masked sub-classifications focused on the F_o_ stator identified a series of intermediates that provided detailed information on conformational substates sampled by the complex during the enzyme cycle (Fig. S2).

**Figure 1:**
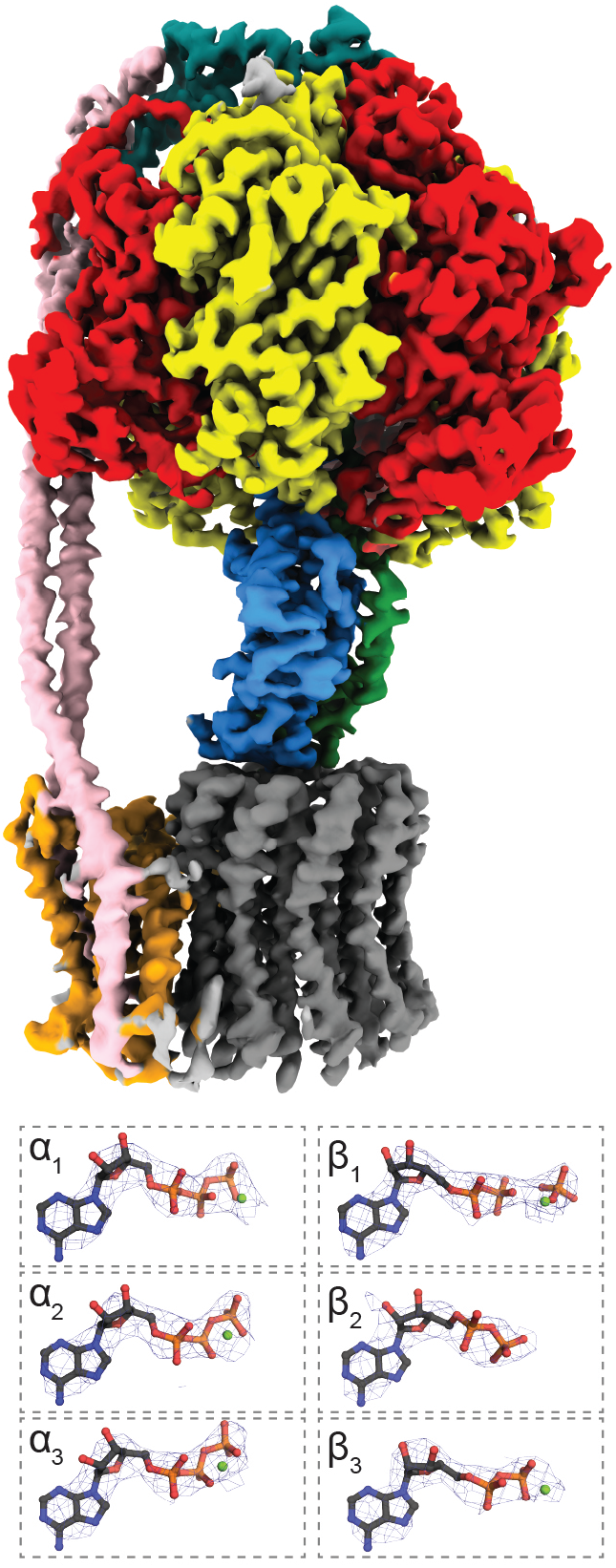
Cryo-EM structure of *E. coli* F_1_F_o_ ATP synthase. *Above:* Cryo-EM volume of *E. coli* F_1_F_o_ ATP synthase State 1a (see Fig. S2 - which shows the best overall detail) rendered and coloured using ChimeraX (*33*). Subunit *a* in red, β in yellow, γ in blue, ε in green, δ in teal, *a* in orange, *b* in pink and *c* in grey. *Below:* Nucleotide occupancy of the *a* and β subunits in State 1a labelled as in (*31*), with corresponding cryo-EM density shown as a blue wire mesh.

The previously identified major conformational states (termed “State 1”, “State 2” and “State 3”) of the enzyme, where the central stalk is rotated by ~120° between each state (*16*), were also identified in the current study with resolutions of 3.1, 3.4 and 3.2 Å, respectively (Fig S2). Masked classification focused on the F_o_ stator of rotational State 1 highlighted four subclasses that describe more subtle movements within the complex (Fig. 2, S2 and S3) and which produced maps of the intact complex that showed much clearer density for the F_o_ region (Fig. S2). The defining theme of these subclasses was flexing of the central and peripheral stalks, with a bending at “pivot 1”, a twist at “pivot 2” and a swivel at “pivot 3” (Fig. S3 and Movie S1). Independently these movements do not facilitate rotation of the *c*_10_ ring (as seen in states 1b-d, Fig. S3). However, in combination (as seen by comparing states 1a and d) they can to accommodate a ~35° rotation of subunits α_3_β_3_δγε*c*_10_ (corresponding to the F_1_ motor plus the *c*-ring) relative to the stator (Fig. 2 and Movie S2). This rotation corresponds to a single *c*-ring sub-step in the F_o_ motor which we propose is driven by thermal fluctuations within the *E. coli* F_1_F_o_ ATP synthase complex, consistent with a Brownian ratchet mechanism. Because these data were obtained at 22°C, they point to the sub-step rotation of the *c*-ring being within a relatively shallow energy landscape, resulting in a highly efficient enzyme. Moreover, because we did not observe any maps corresponding to rotations of a *c*-ring less than a single sub-step, there must be a small energy barrier in between the two *c*-ring positions observed, likely due to the bridge between *a*Arg210 and *c*Asp61 holding the *c*-ring in a energy minimum (*18*). The overall flexibility of the peripheral stalk in *E. coli* ATP synthase observed in the present study is consistent with its predicted mechanical properties that derive from its being constructed from a parallel right-handed coiled coil (*14, 19*) and enable its bending and twisting to facilitate a fluid coupling between stator and the rotor as it rotates. In addition, the movements we observe here may account for the recoil rotation observed using single molecule studies on *E. coli* F_1_F_o_ ATP synthase (*10*), where a small counter-rotation was seen after each sub-step of the F_o_ motor.

**Figure 2:**
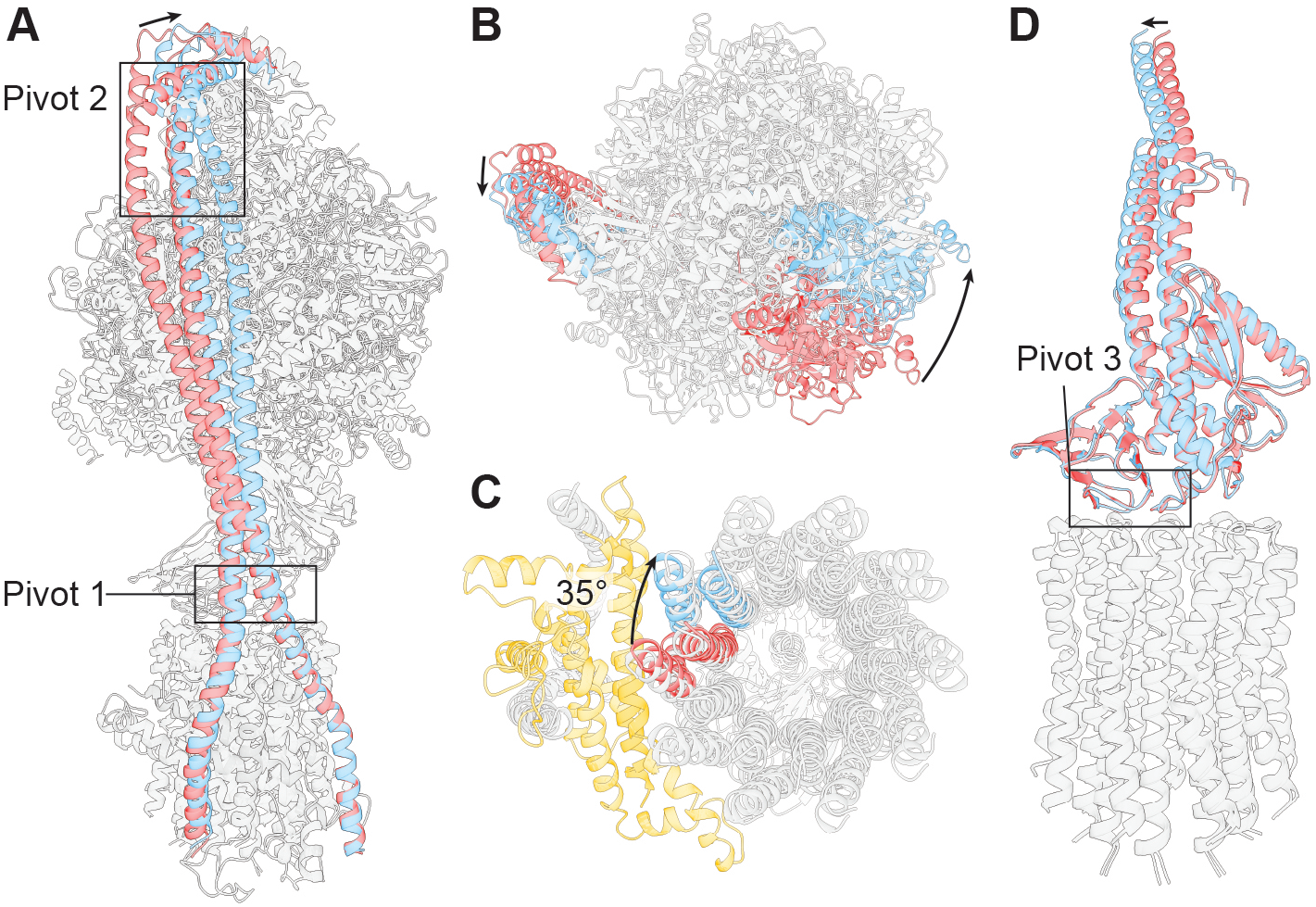
The peripheral stalk bends and twists allowing the *c*-ring to rotate. (**A**) Molecular models of States 1a and 1d (see Fig. S2) superposed on stator subunit *a*. Peripheral stalk of State 1a colored in red and State 1d in cyan. Two pivot points (labelled with black boxes) are required to adapt to the rotational sub-step. (**B**) View from above highlights rotational movement of the of the α_3_β_3_γεδ*c*_10_ relative to the stator (black arrows). (**C**) View from below shows the 35° rotation of α_3_β_3_γεδ*c*_10_ relative to the *a* subunit (shown in orange). (**D**) Molecular models of the central rotor of States 1a and 1d superposed on the c-ring highlights a third pivot at the interface between the stalk and *c*-ring.

In our data on the *E. coli* enzyme we did not observe any hinge movement in the δ subunit, which corresponds to *OSCP* that has been proposed (*15*) to provide flexibility in mitochondrial ATP synthases (Fig. S4). In contrast to *Polytomella* mt. ATP synthase, where the peripheral stalk remained essentially unchanged during rotation (*15*), the *E. coli* peripheral stalk bends and twists substantially. This difference could derive from *Polytomella* mt. ATP synthase having a large bridged peripheral stalk, the structure of which appears likely to inhibit substantial internal movement within this component (*15*) so that, in this case, flexibility is generated primarily by the junction between it and the F_1_ motor, formed by the *OSCP* subunit. However, it is thought (*15*) that this hinge is also found in bovine mitochondrial ATP synthase (*20*), raising the possibility that the mechanism by which the stator flexes might be different between eukaryotes and prokaryotes. Although further work will be needed to investigate this possibility, it holds out the possibility that the hinges present in the *E. coli* stator could represent a novel drug target, especially since a number of agents are known to target the flexing *OSCP* subunit in mitochondrial ATP synthases (*15*). An alternative hypothesis is that the two mechanisms; pivoting in the peripheral and central stalks (seen in this study on *E. coli*), and a hinge in subunit δ/*OSCP* (seen in *Polytomella (15)*), could combine to allow a fluid motion across a ~60° sub-step.

The subclassified maps also provided substantially improved detail in the F_o_ region, compared to that seen without masked classification or that seen previously (*16, 17*). As a result, although their overall resolution was lower than those from the complete ensemble, they enabled a detailed model of the membrane embedded *c, a* and *b* subunits to be built (Fig. 3A). The model generated complements those that have been suggested for other species, particularly for the related PS3 enzyme (*21*), and importantly, because *E. coli* F_1_F_o_ ATP synthase has been a model system for studying ATP synthase for decades, it provides a crucial background to understand the wealth of mutagenesis studies that have been performed in this system. For example, the structure generated for *E. coli* indicates why *a*Arg210 (*22*) is essential for F_o_ rotation because it is found to bind to proton translocating *c*Asp-61 (*23*) (Fig. 3B), along with residues *a*Ser206 (*24*) and *a*Asn214 (*25*) which are seen in a flanking position likely forming the two proton half-channels (*18*) (Fig. 3A). Our model also places *a*Gln252 adjacent to *a*Arg210, which if substituted for one another can recover function (Fig. 3A) (*26*). Although a coordinated metal ion has been proposed to mediate protonation of the *c* subunit (*15*) in related ATP synthases (*15, 27, 28*), close inspection of our maps (Fig. S5) failed to show non-peptide density in this region, suggesting that either our maps do not contain enough detail to identify the metal ion, or it was removed during the purification process, or it is not conserved in this system. Indeed, *a*His248 and *a*His252 in *Polytomella* mt. ATP synthase, which coordinate the ion, correspond to *a*Glu219 and *a*Iso223 in the *E. coli* enzyme, which may explain why we do not observe a metal ion at this position in *E. coli* ATP synthase. Nearby residue *a*His245 in *E. coli* (*a*Glu288 in *Polytomella*) is known to be essential for function (*24*), but inspection of our map in this area again did not lead to any non-peptide density being discovered.

**Figure 3:**
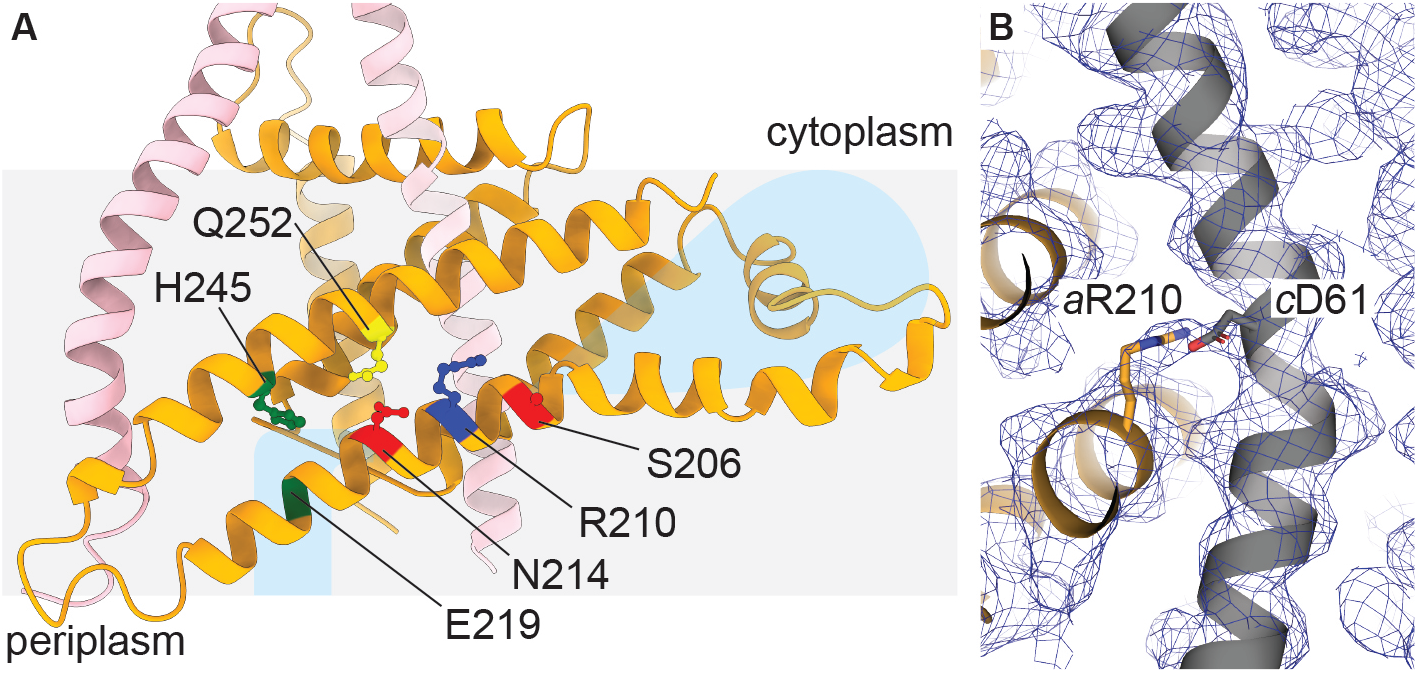
The F_o_ stator of *E. coli* F_1_F_o_ ATP synthase. (**A**) Molecular model of the F_o_ stator, as viewed from the *c*-ring, showing important residues at the stator rotor interface. Colors as in Fig. 1, with membrane region depicted with a grey box and proton half channels shown as light blue regions. *a*Arg210 (dark blue) (*22*) is located adjacent to *c*Asp61 (close up with corresponding EM map [State 1c] shown in **B**). *a*Gln252 is situated proximally to *a*Arg210, which when substituted for one another can regain function (*26*). *a*Asn214 (*25*) defines the beginning of the periplasmic proton channel, and *a*His245 (*24*) and *a*Glu219 (*34*), which are essential for proton flow, are situated on the path out of the membrane. *a*Ser206 (*24*) defines the beginning of the cytoplasmic proton channel.

The cryo-EM structure of the F_1_ motor described here also provided information on the structural changes and nucleotide occupancy introduced by the binding of MgADP. To date, only two studies have provided cryo-EM maps of an ATP synthase in which conditions have been manipulated to image the sample with added nucleotide (*17, 29*). One of these studies, which used a short incubation with MgATP and *E. coli* enzyme, described the molecular events that occur upon ATP binding and showed that the C-terminal domain of the inhibitory subunit ε (*30*) was removed from the central cavity when ATP was bound and this, in turn, allowed rotation of the rotor (*17*). Our present maps show that incubation of 10 mM MgADP induces the catalytic β subunits to bind nucleotide (Fig. 1) which, in turn, induces a partial “closure” of the subunit that is blocked in a “half-open” state by the inhibitory ε subunit in a manner reminiscent of that seen the crystal structure of isolated *E. coli* F_1_-ATPase (*31*). This strongly suggests that a key function of the ε subunit is to increase the efficiency of the enzyme by preventing *E. coli* F_1_F_o_ ATP synthase from entering the low energy MgADP inhibited state, which is known to inactivate the enzyme (*32*) (Movie S3).

The work presented here has generated a comprehensive structural model of *E. coli* F_1_F_o_ ATP synthase providing a framework to understand mutagenesis studies together with yielding insight into the flexibility of the peripheral and central stalks. The substates observed suggesting an elegant mechanism by which the F_1_ and F_o_ motors can be coupled with minimal energy loss and, in addition, the structural rearrangement observed on binding of MgADP indicates that the ε subunit can function as a brace to prevent the complex entering the MgADP inhibited state.

## Acknowledgments

We wish to thank and acknowledge Dr Craig Yoshioka and Dr Claudia López (Oregon Health & Sciences University (OHSU)) for data collection and processing expertise; Dr Thomas Duncan (Department of Biochemistry & Molecular Biology, SUNY Upstate Medical University, Syracuse, NY, USA) for initial discussions on experimental design; and Yi Cheng Zeng (the Victor Chang Cardiac Research Institute, NSW, Australia) for aiding in particle picking. A.G.S was supported by a National Health and Medical Research Council Fellowship APP1159347 and Grant APP1146403. We wish to thank and acknowledge the use of the Victor Chang Innovation Centre, funded by the NSW Government, and the Electron Microscope Unit at UNSW Sydney, funded in part by the NSW Government. A portion of this research was supported by NIH grant U24GM129547 and performed at the Pacific Northwest Centre for Cryo-EM at OHSU and accessed through EMSL (grid.436923.9), a DOE Office of Science User Facility sponsored by the Office of Biological and Environmental Research.; Author contributions: A.G.S. conceived and supervised the study, and wrote the original draft. M.S. performed the formal analysis of the study (purified protein, prepared cryo-EM grids, performed reconstructions). J.W. built the atomic models into the cyro-EM maps. R.I. edited the manuscript and conceived the study; Data was deposited in the EMDB and PDB with the following accession codes: State 1a; EMD-20167 & PDB-6OQR. State 1b; EMD-20168 & PDB-6OQS. State 1c; EMD-20169 & PDB-6OQT. State 1d; EMD-20170 & PDB-6OQU. State 2a; EMD-20171 & PDB-6OQV. State 3a; EMD-20172 & PDB-6OQW.

