## Supplemental methods, figures and table for "Visualizing movements in *E. coli* F_1_F_o_ ATP synthase indicates how the F_1_ and F_o_ motors are coupled"

### Materials and Methods

Protein purification

The *E. coli* $F_1F_o$ ATP synthase protein was prepared as described in Sobti et al. 2019 *(17)*. Cysteine-free *E. coli* ATP synthase (all cysteines residues substituted with alanine and a His-tag introduced on the β subunit) was expressed in *E. coli* DK8 strain (*35*). Cells were grown at 37°C in LB medium supplemented with 100 µg/ml ampicillin for 5 h. The cells were harvested by centrifugation at 5,000 g, providing ~1.25 g cells per litre of culture. Cells were resuspended in lysis buffer containing 50 mM Tris/Cl pH 8.0, 100 mM NaCl, 5 mM $MgCl_2$, 0.1 mM EDTA, 2.5% glycerol and 1 µg/ml DNase I, and processed with three freeze thaw cycles followed by one pass through a continuous flow cell disruptor at 20 kPSI. Cellular debris was removed by centrifuging at 7,700 × g for 15 mins, and the membranes were collected by ultracentrifugation at 100,000 × g for 1 h. The ATP synthase complex was extracted from membranes at 4°C for 1 hr by resuspending the pellet in extraction buffer consisting of 20 mM Tris/Cl, pH 8.0, 300 mM NaCl, 2 mM $MgCl_2$, 100 mM sucrose, 20 mM imidazole, 10% glycerol, 4 mM digitonin and EDTA-free protease inhibitor tablets (Roche). Insoluble material was removed by ultracentrifugation at 100,000 g for 30 min. The complex was then purified by binding on Talon resin (Clontech) and eluted in 150 mM imidazole, and further purified with size exclusion chromatography on a 16/60 Superose 6 column equilibrated in a buffer containing 20 mM Tris/Cl pH 8.0, 100 mM NaCl, 1 mM digitonin and 2 mM $MgCl_2$. The purified protein was then concentrated to 11 µM (6 mg/ml), and snap frozen and stored for grid preparation.

Cryo-EM grid preparation

1 µl of 100 mM ADP/100 mM $MgCl_2$ (pH 8.0) was added to an aliquot of 9 µl of purified cysteine-free *E. coli* $F_1F_o$ ATP synthase at 11 µM (6 mg/ml) and the sample was incubated at 22 °C for 30 s, before 3.5 µl was placed on glow-discharged holey gold grid (Ultrafoils R1.2/1.3, 200 Mesh). Grids were blotted for 3 s at 22 °C, 100 % humidity and flash-frozen in liquid ethane using a FEI Vitrobot Mark IV (total time for sample application, blotting and freezing was 15 s).

Data collection

Grids were transferred to a Thermo Fisher Talos Arctica transmission electron microscope (TEM) operating at 200 kV and screened for ice thickness and particle density. Grids were subsequently transferred to a Thermo Fisher Titan Krios TEM operating at 300 kV equipped with a Gatan BioQuantum energy filter and K3 Camera at the Pacific Northwest Centre for Cryo-EM at OHSU. Images were recorded automatically using serial EM at 81,000 x magnification yielding a pixel size of 0.54 Å (K3 operating in super resolution mode). A total dose of 48 electrons per $Å^2$ was used spread over 77 frames, with a total exposure time of 3.5 s. 9,342 movie micrographs were collected (Figure S1).

Data processing

MotionCorr2 (*36*) was used to correct local beam-induced motion and to align resulting frames, with 3x3 patches and binning by a factor of two. Defocus and astigmatism values were estimated using Gctf (*37*) and 8,290 micrographs were selected after exclusion based on ice contamination, drift and astigmatism. 1,361 particles were manually picked and subjected to 2D classification to generate templates for autopicking in RELION-3.0 (*38*), yielding 1,349,270 particles. Images were manually inspected to unpick regions containing ice or aggregated protein to yield 1,111,931 particles. These particles were binned by a factor of four and subjected to 2D classification generating a final dataset of 709,190 particles. These particles when then re-extracted at full resolution and further classified into 3D classes using a low pass filtered cryo-EM model generated from a previous study *(16)*, yielding maps related by a rotation of the central stalk (355,964, 179,005 and 174,221 particles). Focussed classification, using a mask encompassing the $F_o$ stator, was implemented without performing image alignment in Relion 3.0, yielding nine defined subclasses (four for State 1, three for State 2 and two for State 3; see Fig. S2 for detailed flowchart describing this classification and S6 for FSC curves). Final refinements were performed in cryoSPARC (*39*).

Model building

Models were built in coot using pdb IDs 3oaa *(31)* (*E. coli* $F_1$-ATPase), 1abv (*40*) (N-terminal domain of *E. coli* subunit δ) and 6n2y *(21)* (Bacillus PS3 ATP synthase) as a guide.

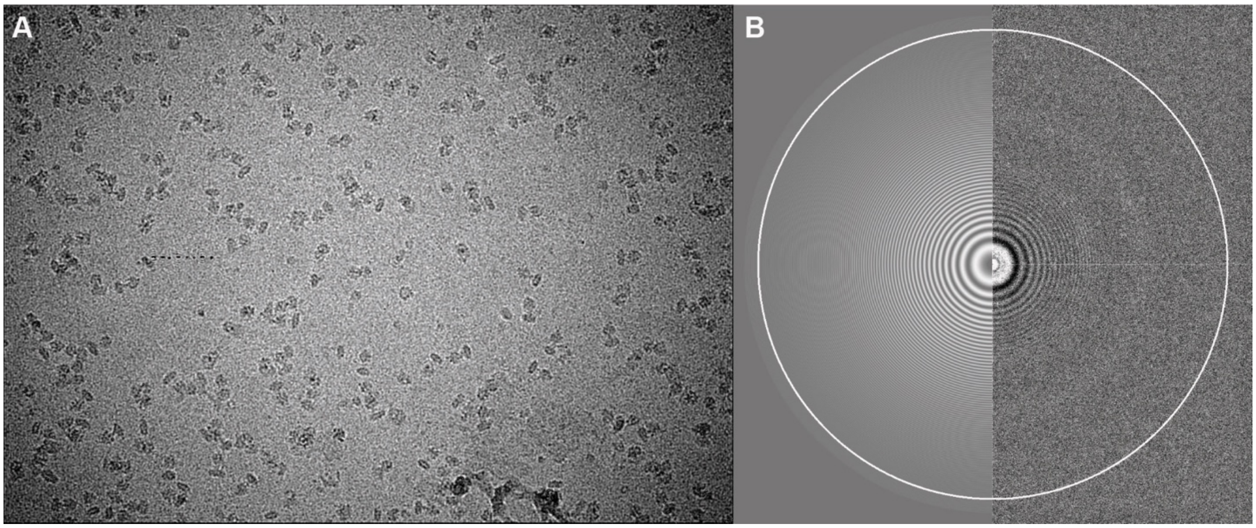


**Fig. S1.**

(**A**) Representative micrograph showing *E. coli* ATP synthase particles and (**B**) Gctf output of same micrograph.

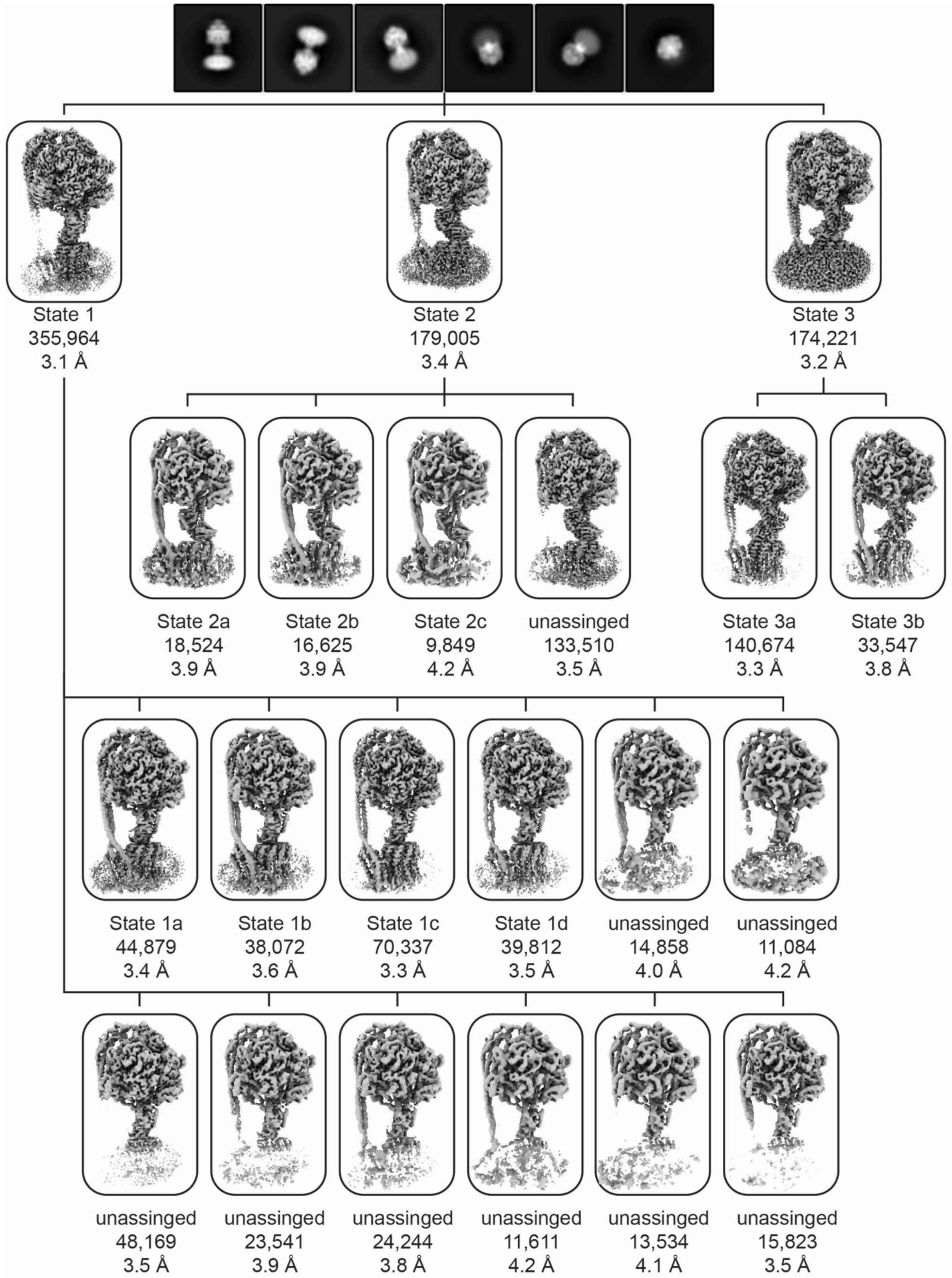


**Fig. S2.**
Flowchart to describe classification of particles into distinct rotary conformations.

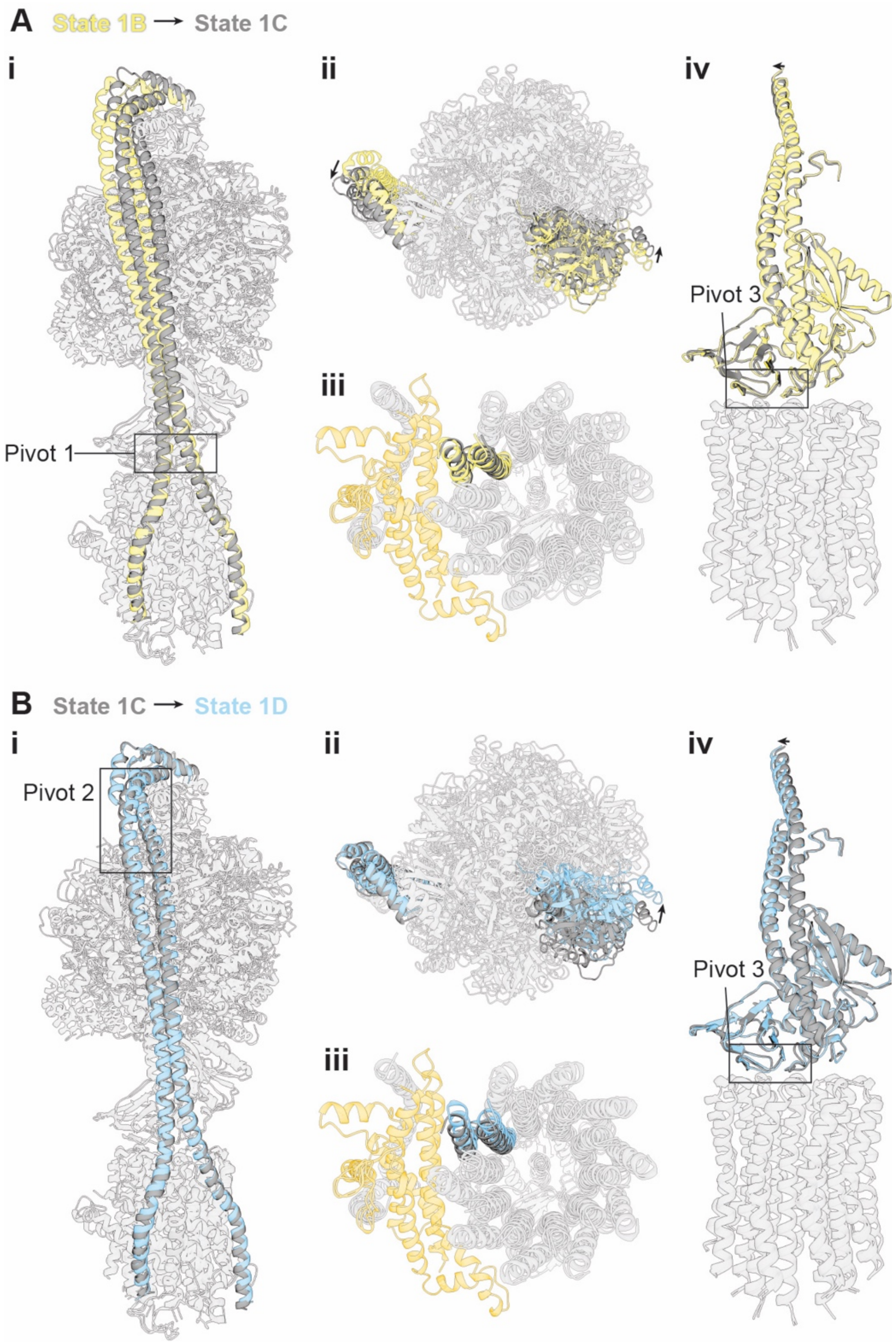


**Fig. S3.**
Comparison of states 1c-d shows pivot points 1 (**A i** and **ii**) and 2 (**B i** and **ii**). Independently, these pivot points do not allow a sub-step (**iv** in **A** and **B**) and pivot 3 (**iii** in **A** and **B**) accommodates this movement.

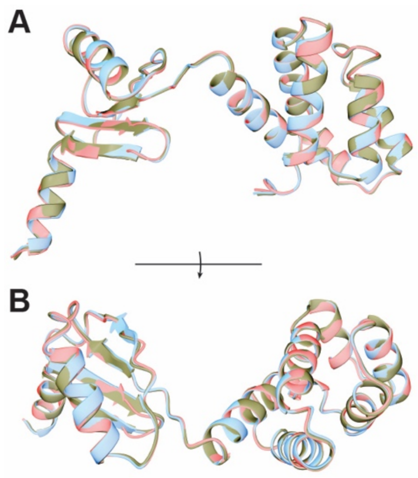


**Fig. S4.**
The structure of subunit δ is very similar between States 1a-d with an r.m.s.d of 0.3 Å between all structures. State 1a in red, 1b in yellow, 1c in black and 1d in blue.

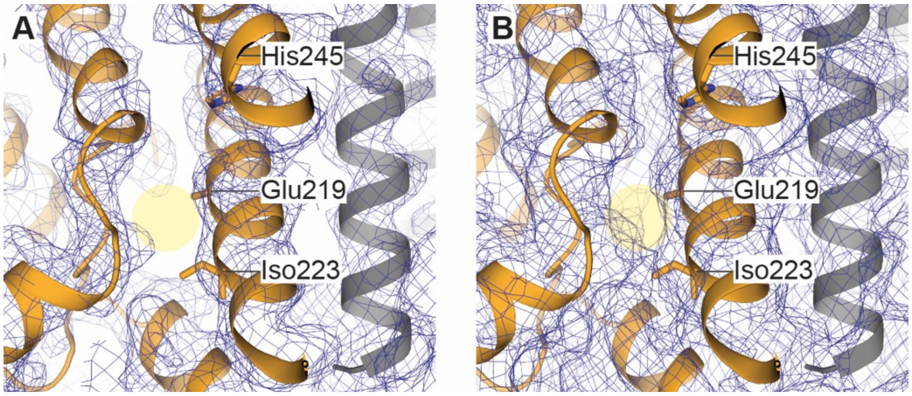


**Fig. S5.**
Close up of putative metal ion binding site. **A** high threshold and **B** low threshold. No non-protein density could be observed in putative metal ion binding site (highlighted in yellow). Glu219, Iso223 and His245 of subunit *a* labelled, with subunit *a* in orange and subunit *c* in grey shown as cartoon.

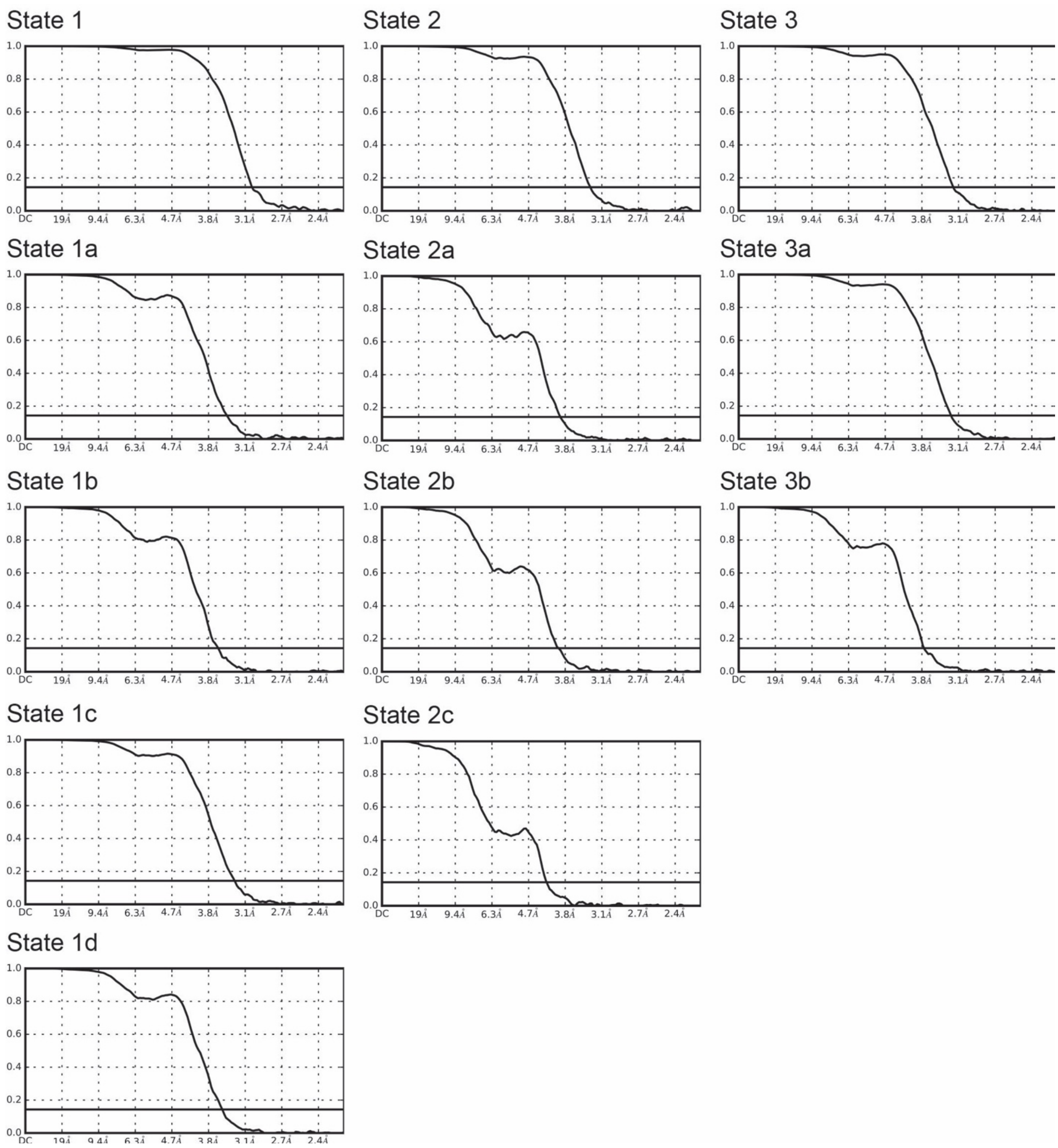


**Fig. S6.**
Fourier shell correlation curves for states shown in Fig. S2.

**Table S1.**
Cryo-EM data processing table.

| Description | EMDB ID | PDB ID | Resolution (Å) | Applied B factor ($Å^2$) | Number of particles |
|---|---|---|---|---|---|
| Substate 1a | 20167 | 6OQR | 3.4 | -80 | 44,879 |
| Substate 1b | 20168 | 6OQS | 3.6 | -80 | 38,072 |
| Substate 1c | 20169 | 6OQT | 3.3 | -91 | 70,337 |
| Substate 1d | 20170 | 6OQU | 3.5 | -76 | 39,812 |
| Substate 2a | 20171 | 6OQV | 3.9 | -65 | 18,524 |
| Substate 3a | 20172 | 6OQW | 3.3 | -102 | 140,674 |

**Movie Captions:**

**Movie S1:** Movie to show interpolation between States 1b, 1c and 1d of *E. coli* $F_1F_o$ ATP synthase (see Fig. S2 for details). Flexing and bending can be observed in the central and peripheral stalks. **A** and **B**; side views. C top view.

**Movie S2:** Movie to show interpolation between State 1a and 1d of *E. coli* $F_1F_o$ ATP synthase (see Fig. S3 for details). The peripheral stalk bends and twists to accommodate a rational sub-step in the $F_o$ motor. **A** and **B**; side views. **C** top view. **D** bottom view.

**Movie S3:** Movie to describe structural changes that occur in *E. coli* $F_1F_o$ ATP synthase upon incubation of ADP and ATP. The movie starts with the conformation of the complex incubated with 10 mM MgADP (this study), which shows a β subunit (yellow) in a "half-closed" position, bound to the ε C-terminal domain (green - in an "up" position) which is preventing the complex from entering the ADP inhibited state. Next, without added nucleotide, the β subunit transitions to an "open" conformation (as seen in *(16)* - imaged without addition of nucleotide) and the ε C-terminal domain can escape from the $F_1$ motor. The ε C-terminal domain transitions from the "up" to the "down" position and the β subunit transitions to a "closed" active position (as seen in *(17)* - imaged after the addition 10 mM MgATP).
